# Detection and full genome sequencing of a Deltacoronavirus and other bird associated viruses from feces of the kelp gull (*Larus dominicanus*) sampled at the South Shetland Islands Antarctica

**DOI:** 10.1101/2024.10.14.618326

**Authors:** Fernanda Gomes, Alexandre Freitas da Silva, Tatiana Prado, Paola Cristina Resende, Leonardo Corrêa Da Silva, Wim Degrave, Maithê Magalhães, Adriana Vivioni, Yago José Mariz Dias, Marilda Siqueira, Martha Brandão, Luciana R. Appolinario, Gabriel da Luz Wallau, Maria Ogrzewalska

## Abstract

Bird species are known to be the main reservoir of a range of respiratory viruses such as Influenza, Newcastle and Coronaviruses. Migratory birds are particularly important for the maintenance and long distance spread of the virus to wild bird and poultry species but eventually to mammal species as well. Antarctica’s pristine environment and wildlife is of immense biological value, but the spread of such deadly viruses pose a substantial threat to the region’s fragile ecosystems. To investigate the presence of respiratory viruses in the region we sampled feces of different wild migratory birds at various localities in the South Shetland Islands in the Antarctic summer of 2023 and screened them for coronaviruses (CoVs) and influenza A virus (IAV). Viral screening was performed by the conventional pancoronavirus RT-PCR protocol (CoVs), by quantitative one-step real-time RT-PCR (IAVs) followed by metatranscriptomic sequencing of positive samples. During January and February of 2023, we collected and examined a total of 243 fecal samples representing *Stercorarius* spp (N=5), *Larus dominicanus* (N=16), *Phalacrocorax bransfieldensis* (N=3), *Pygoscelis adeliae* (N=19), *Pygoscelis antarcticus* (N=38), *Pygoscelis papua* (N=139), *Pygoscelis* spp (N=23). All tested samples were negative for influenza A and one sample from the colony of *L. dominicanus* at Keller Peninsula, King George Island, tested positive for CoVs. Metatranscriptomic sequencing recovered a full deltaCoV genome. Nucleotide and amino acid distance analysis revealed that the deltacoronavirus detected belongs to subgenus *Buldecovirus* and to the novel wild bird deltaCoV clade previously identified infecting Antarctica penguins. The identified deltaCov is most closely related to a deltacoronavirus previously identified 2014 in *P. papua* penguin sampled at Isla Kopaitik, Base O’Higgin suggesting a potential cross species transmission. The presence of CoVs in Antarctic migratory seabirds raises concerns about their impact on the wild bird population in Antarctica and their potential role in virus dispersion through intra and intercontinental migratory routes. These findings contribute valuable insights into virus dynamics among seabird populations, laying the groundwork for future investigations in this field and warning of the importance of viral surveillance on the Antarctic fauna.

**Repositories:** The raw reads were submitted to NCBI Sequence Read Archive (SRA) database and are available under project number: PRJNA1160912 and Run Accessions: SRR30664529 and SRR30664530.

**Impact statement:** This study provides insights into the presence of respiratory viruses, specifically coronaviruses (CoVs), in Antarctic seabirds. By detecting a novel wild bird deltacoronavirus in gulls’ population in the South Shetland Islands, our research contributes to the growing body of literature on viral transmission in remote ecosystems. The findings highlight the potential of migratory birds to act as reservoirs and vectors of viruses in the Antarctic region and underscores the urgent need for continued viral surveillance.

## INTRODUCTION

Antarctica harbors a rich and unique endemic fauna, including several bird species (penguins, petrels, cormorants, gulls, skuas terns), along with its associated microbiome (Shirihai, 2008). The geographic isolation of this continent coupled with the extreme climate and unique ecological interactions has historically been assumed to protect the indigenous Antarctic wildlife from exposure to infectious agents (Convey and Stevens, 2007). However, this pristine environment has been threatened in recent decades, especially because of increased presence and human activities that are compromising the ecosystem health such as tourism, marine exploitation and increase of waste disposal (Convey and Peck, 2019; Hwengwere et al., 2022; Woehler et al., 2014). The invasion of exotic or non-native species coupled with climate change are additional factors that contributes to ecological changes in different ways including the introduction of new wildlife pathogen and potential population size reduction of endangered species due to mortality (Banyard et al., 2024; Barbosa et al., 2020; Woehler et al., 2014). Thus, surveillance approaches that address animal and human health under a One Health framework are key to characterize and mitigate the impact of pathogenic viruses circulating in Antarctica wildlife (Convey and Peck, 2019; Nath and Sindwani, 2023).

Seabirds are infected by a variety of microorganisms including protozoa, bacteria, and viruses (Coffee et al., 2010; Lee et al., 2014; McCoy et al., 2016; Park et al., 2012). Although previous studies have demonstrated the presence of a variety of viruses affecting the Antarctic avifauna (Hurt et al., 2016; Hurt et al., 2014; Smeele et al., 2018; Wille et al., 2019; Zamora et al., 2023) our comprehension of the endemic viral diversity in this region, which includes potentially zoonotic pathogens remains limited. Among viruses infecting seabirds, influenza avian viruses (IAVs) and coronaviruses (CoVs) warrant special attention due to their potential to cause high mortality in both wild and domestic birds and hence inducing significant impacts on these populations (Banyard et al., 2024; Chu et al., 2011; Muradrasoli et al., 2010).

Coronaviruses belong to the *Coronaviridae* family, within the *Nidovirales* order (Cui et al., 2019) and infect a range of animal species. They are positive-sense RNA viruses classified into four genera based on their phylogenetic relationships: Alphacoronavirus, Betacoronavirus, Gammacoronavirus, and Deltacoronavirus. Alpha and beta coronaviruses primarily infect mammals, while gamma and delta coronaviruses primarily infect birds, although mammals infection has also been reported (Cui et al., 2019). Despite coronaviruses being known to circulate in wildlife populations there is limited investigation of coronaviruses in Antarctic fauna. In the absence of more extensive monitoring programs, it is not possible to determine which viral species circulate; which animals are potential reservoirs; what is the prevalence or burden of these viruses in animal hosts and the spillover risks (Barbosa et al., 2020).

Different subtypes of influenza A viruses that infect humans and animals have the potential to cause large epidemics making these viruses an important animal health concern (Uyeki et al., 2022). Influenza A viruses are negative-sense single-stranded segmented RNA viruses belonging to the *Orthomyxoviridae* family. Influenza A viruses in humans originated from birds and swine and currently different subtypes that infect wild birds can infect humans (Newman et al., 2008; Uyeki et al., 2022). Wild birds (mainly Anseriformes and Charadriiformes) are recognized as the natural reservoir of avian influenza viruses (AIVs) (Olsen et al., 2006; Webster et al., 1992). It is estimated that 100 million seabirds live in the Antarctic Peninsula and adjacent islands, regularly encountering migratory birds that use the islands for nesting (de Seixas et al., 2022; Shirihai, 2008). The ecology and migratory patterns of these birds probably have a direct effect on the global distribution and diversity of AIVs (Hurt et al., 2016; Hurt et al., 2014). Especially concerning is the recent spread of highly pathogenic influenza A subtype of the H5N1 strain to the Antarctic continent (Banyard et al., 2024; Dewar et al., 2022).

To fill the surveillance gaps about CoVs and AIVs virus in Antarctic wildlife we performed targeted viral surveillance for coronavirus and influenza A virus in fresh fecal samples from various Antarctic bird species sampled at South Shetland Archipelago during the austral summer 2023. We detected a single fecal sample positive for a deltacoronavirus and recovered its full genome. This virus belongs to the *Buldecovirus* subgenus and is clustered with a full genome from a deltacoronavirus circulating in the same region detected in the Gentoo Penguin (*Pygoscelis papua*) sampled in 2014 supporting long-term maintenance of the virus in the region and a potential cross-species transmission event.

## MATERIALS AND METHODS

### Sample collection

Fecal samples were collected during field expeditions conducted in the South Shetland Islands, near the Antarctic Peninsula, in January and February 2023. The sampling efforts covered eight islands and 14 different localities (**Figure 1**). During the sampling process, we collected individual fresh samples from monitored animals as well as fecal material from penguins’ nesting sites. Fecal samples were obtained using sterile Dacron swabs, which were immediately placed into tubes containing 1 mL of Viral Transport Medium, following the established protocol (Ogrzewalska et al., 2022). Subsequently, the samples were refrigerated for a maximum of 4 hours before being frozen at -80°C for subsequent analysis.

**Figure 1.**
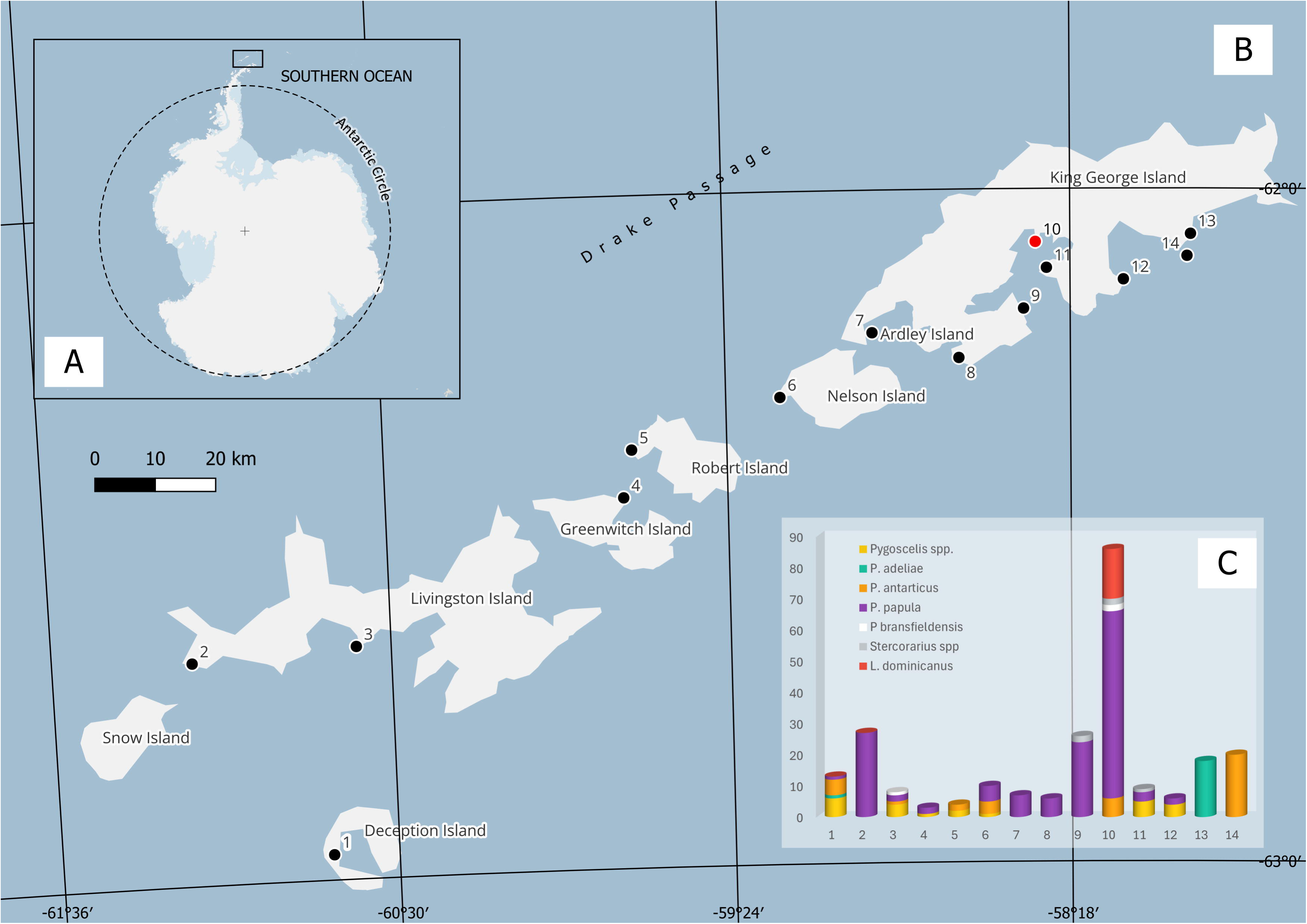
A) Localization of the South Shetland Islands collection sites in the present study, B) Localization of the sampling sites along South Shetland Islands in January-February 2023. 1 - Fumarole Bay, 2- Byers Peninsula, 3 - Hannah Point, 4 - Maldonado Base, 5- Coppermine Peninsula, 6 - Harmony Point, 7 - Ardley Island, 8 - Potter Peninsula, 9 – Copacabana, 10 - Keller Peninsula, 11 - Point Hennequin, 12 - Lions Rump, 13 - Three Sister’s Point. In red marked where the novel coronavirus was found. C) Number of samples collected form each sampling area per species. More details in Supplementary Table 1. Map made using a package from the platform Quantarctica 3.2 database in the software QGIS 3.32.2.

### Viral RNA extraction

Clarified fecal suspensions (20% wt/vol) were prepared as previously described (Ogrzewalska et al., 2022). Subsequently, 140 mL of the supernatant was utilized for viral RNA extraction. The extraction process was conducted using a QIAamp viral RNA minikit (Qiagen, CA, USA) and a QIAcube automated system (Qiagen), following the manufacturer’s guidelines. The isolated RNA was promptly stored at -80°C until further molecular analysis. Negative controls for each extraction procedure were carried out using RNase/DNase-free water.

### Coronavirus (CoVs) screening by conventional pancoronavirus RT-PCR

All the samples were also subjected also to pancoronavirus PCR targeting the RNA-dependent RNA polymerase (*RdRp*) gene as described previously (Chu et al., 2011; Gomes et al., 2023; Woo et al., 2005). In brief, RNA was first amplified using a first-round PCR (RdRp S1 5ߣ-GGKTGGGAYTAYCCKAARTG -3’, RdRp R1 5’-TGYTGTSWRCARAAYTCRTG-3’) with the One-Step RT-PCR Enzyme MixKit (Qiagen), targeting a total expected size of 620 base pairs (bp). Subsequently, a second PCR was performed using the Phusion RT-PCR Enzyme Mix kit (Sigma-Aldrich) and primers Bat1F 5’- GGTTGGGACTATCCTAAGTGTGA -3’ and Bat1R 5’-CCATCATCAGATAGAATCATCAT-3’, with 1 μL of the first-round amplified product as template. The resulting amplicons (∼440 bp) were visualized on 1.5% agarose gels stained with SYBR™ Safe DNA Gel Stain (Thermo Fisher Scientific). For Sanger sequencing, DNA was purified using the QIAquick Gel Extraction Kit (Qiagen) following the manufacturer’s recommendations. The Sanger sequencing reaction was prepared utilizing the BigDye Terminator v3.1 Cycle Sequencing Kit (Life Technologies) with primers Bat1F and Bat1R at a concentration of 3.2 pmoles. Sequencing was executed on the ABI 3730 DNA Analyzer (Applied Biosystems) following the protocols established by (Otto et al., 2008). The final consensus sequences were derived using Sequencher 5.1.

### Influenza A virus (IAV) screening by quantitative one-step real-time RT-PCR

The screening for IAVs was conducted utilizing the commercial RT-PCR kit (Biomanguinhos, Fiocruz). Samples exhibiting a characteristic sigmoid curve and crossing the threshold line below a threshold cycle (CT) value of 38 were considered positive.

### Metatranscriptomic sequencing

After validation of a deltacoronavirus detection in the F1596 sample by Sanger sequencing, we processed the same sample by metatranscriptomics with the goal to recover the full deltacoronavirus genome. The positive sample was treated with Ambion® TURBO DNA-free™ Kit (Invitrogen) to remove residual genomic DNA from the extracted RNA. Initially, 30 µL of RNA was combined with 3 µL of 10x TURBO DNase Buffer and 1 µL of TURBO DNase in a microcentrifuge tube. The mixture was incubated at 37°C for 30 minutes to allow the DNase to degrade any contaminating DNA. Following this incubation, 3 µL of Stop Solution was added, and the sample was further incubated for 5 minutes at room temperature to ensure complete inactivation of the DNase. The sample was then centrifuged at 10,000 × g for 1.5 minutes, and the supernatant was transferred to a new tube. To deplete the host rRNA we used the Illumina Ribo-Zero Plus rRNA Depletion Kit (Illumina, San Diego, CA, USA). The kit employs capture oligonucleotides and hybridization techniques to eliminate rRNA, thereby enriching messenger RNA (mRNA) and other types of RNA of interest. After rRNA depletion, the remaining RNA is converted into cDNA and prepared for library construction using the Illumina DNA Prep Kit, which includes cDNA fragmentation and the addition of necessary adapters for sequencing on Illumina platforms. The libraries were sequenced on the Illumina® NextSeq platform (Illumina, San Diego, CA, USA) using a NextSeq 1000/2000 P2 cartridge. The raw reads were submitted to NCBI Sequence Read Archive (SRA) database and are available under project number: PRJNA1160912 and Run Accessions: SRR30664529 and SRR30664530.

### Deltacoronavirus genome characterization and phylogenetic analysis

Raw reads generated from Nextseq Illumina sequencing were first analyzed on fastp v0.23.2 to trim low-quality reads with a cutoff of 20 for Phred score and a minimum read length of 36 bp. Trimmed reads were used to perform a *de novo* assembly analysis on metaSPAdes v3.15.5 in default mode (Nurk et al., 2017). All contigs were submitted to a Diamond blastx analysis (Buchfink et al., 2015) against a custom dataset composed of all viral proteins assigned to taxonomy tag (txid10239) on NCBI plus RdRp sequence databases such as NeoRdRp (Sakaguchi et al., 2022), PalmDB (Edgar et al., 2022) and RdRp-Scan (Charon et al., 2022) as performed previously by (da Silva et al., 2024). The sequences showing the best hits on viral sequences were submitted to a second Diamond blastx analysis against the non- redundant (NR) database from NCBI to remove false negative hits. Additional BLASTn (Altschul et al., 1990) analyses were conducted against the NT database from NCBI. Only contigs showing hits against viral proteins in these two analyses were kept and further analyzed. Contigs annotated as viral were analyzed on ViralComplete (Antipov et al., 2020) tool to classify them into full-length or partial according to their completeness in relation to NCBI RefSeq genomes. The trimmed reads were then mapped against the contigs using Bowtie2 in default mode. The BAM file was sorted and indexed by Samtools 1.20 (Danecek et al., 2021) and genome metrics were calculated using the coverM tool (https://github.com/wwood/CoverM). Genomic maps were constructed using the BAMdash tool (https://github.com/jonas-fuchs/BAMdash) and *gggenomes* R package (https://github.com/thackl/gggenomes).

For the phylogenetic analysis, we downloaded all closely related deltacoronavirus sequences from NCBI virus using the F1596 deltacoronavirus as a query from a blastn analysis implemented within the database (https://blast.ncbi.nlm.nih.gov/Blast.cgi). All deltacoronavirus RdRp sequences from NCBI virus were also included in our analysis. Additionally, we included reference sequences from *Alpha* (OQ540913.1)*, Beta* (LC706864.1) and *Gamma* (MN509588.1) coronaviruses to root our trees. All retrieved sequences were aligned together with the sequenced genome using MAFFT v7.511. Nucleotide alignments were inspected for recombination using SplitsTree App v.6.3.35 running the Pairwise Homoplasy Index (Phi) test (Bruen et al., 2006). For the phylogenetic analysis, we performed three approaches: I- Nucleotide alignment of partial RdRp sequences, II- Nucleotide CDS Alignment of ORF1ab and III - translated CDS amino acid sequences of ORF1ab. The Maximum Likelihood phylogenetic analysis was performed on IQ-TREE 2.3.6 with the SH-aLRT test and ultrafast bootstrap with 1000 replicates. The best evolutionary models were selected by ModelFinder (Kalyaanamoorthy et al., 2017). The trees were visualized and annotated using *ggtree* R package (Yu et al., 2017). Distance matrices were calculated using the alignments in MEGA 11.0.13 performing a p-distance method and rates among sites parameter set as G4+I and gaps/missing data treated with pairwise deletion.

To better understand and visualize potential virus transmission between species, we mapped the distributions of bird species in which closely related coronaviruses were detected (belonging to the new deltacoronavirus clade), using data from the BirdLife database (BirdLife, 2023).

## RESULTS

During the Brazilian Antarctica Expedition in January-February 2023, 243 fresh fecal avian samples were collected from skuas *Stercorarius* spp (N=5), kelp gulls *Larus dominicanus* (N=16), shags *Phalacrocorax bransfieldensis* (N=3), and penguins *Pygoscelis adeliae* (N=19), *Pygoscelis antarcticus* (N=38), *Pygoscelis papua* (N=139), *Pygoscelis* spp (N=23) (Figure 1C, Supplementary Table 1). All tested samples were negative for influenza A and one sample from the colony of *L. dominicanus* at Keller Peninsula, King George Island (**Figure 1B**), tested positive for CoVs (sample F1596). Observed birds did present symptoms indicating any disease.

A total of 64.2 million (32.1 of paired-end) of reads were generated in Illumina sequencing for *L. dominicanus* sample and 52.6 million quality filtered reads were assembled into 7,292 contigs. One contig of 26,093 bp and mean depth coverage of 2240.26x was assembled and identified as a complete genome of a deltacoronavirus. This virus was designated DeltaCoV/AvCoV/Larus_dominicanus/Antarctica/Fiocruz-F1596/2023 (**Figure 2A**). The assembled genome has been deposited in GenBank under PQ351810.1 accession number.

**Figure 2.**
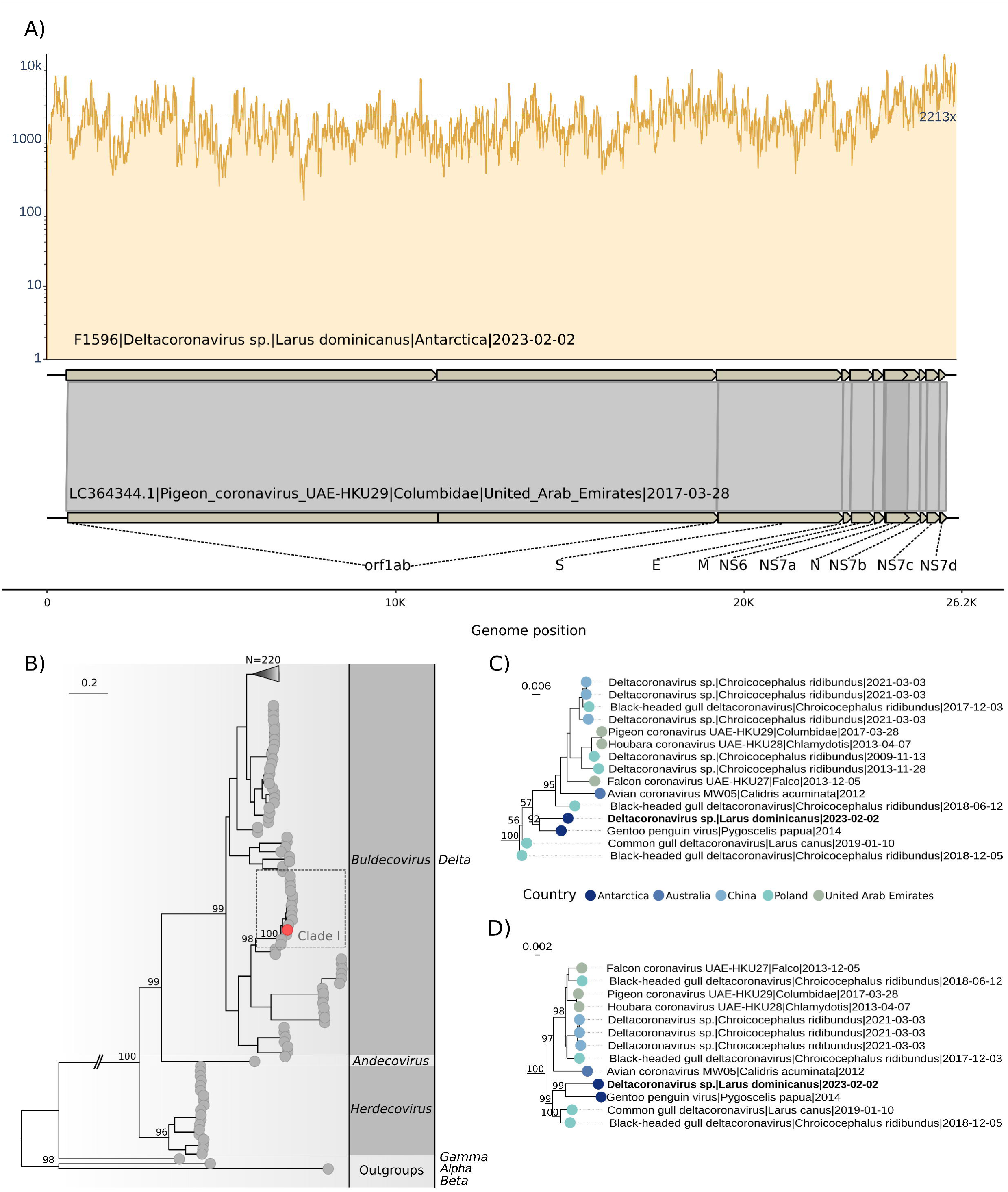
Genomic map and phylogenetic analysis of Deltacoronavirus identified in *Larus dominicanus.* A) Genomic map showing the coverage of Deltacoronavirus F1596 and comparison with the best annotated reference (LC364344.1). B) Phylogenetic analysis of nucleotide RdRp alignment of Deltacoronaviruses using TIM2+F+I+G4 evolutionary model. The outgroups represent sequences from *Alpha* (OQ540913.1)*, Beta* (LC706864.1) and *Gamma* (MN509588.1) coronaviruses. Red tip point shows the positioning of the Deltacov F1596. C) Zoom of clade I. D) Clade I reconstructed based on phylogenetic analysis of amino acid ORF1ab-polyprotein alignment using Q.plant+F+I+G4 evolutionary model. The phylogenetic trees were reconstructed on IQ-TREE 2.3.6 (Nguyen et al., 2015) performing the SH-aLRT test and ultrafast bootstrap with 1000 replicates. Bold tip labels represent the sequence characterized in this study. Tip point colors represent the sampling location.

The Phi test for recombination showed no significant values for recombination on nucleotide alignments. Therefore, three Maximum likelihood phylogenetic analyses were performed based on alignments of RdRp, ORF1ab-CDS and polyprotein encoded by ORF1ab. The RdRp phylogenetic analysis, which included 316 coronaviruses (313 *Delta*, 1 *Alpha*, 1 *Beta* and 1 *Gamma* coronaviruses) positioned the Deltacoronavirus from *L. dominicanus* within the *Buldecovirus* subgenus showed in clade I (Figure 1B). Clade I comprises 15 *Buldecovirus* sequences identified in Antarctica, Australia, China, Poland and United Arab Emirates from other gulls (Figure 1C). *Larus dominicanus Buldecovirus* sequence was closely related with Gentoo penguin virus identified in *P. papua* identified in Antarctica, Isla Kopaitik, Base O’Higgin in 2014 (MT025058.1) with a high node support of 92 in the RdRp phylogeny. This finding was corroborated by the analysis of CDS-ORF1ab (**Supplementary figure 1A**) and polyprotein sequence encoded by ORF1ab, which showed high node supports of 99 and 100, respectively (**Figure 2D**).

The identity percentages between the Deltacoronavirus identified in *L. dominicanus* and other sequences from the NCBI database ranged from 59-94% (median=72.38, stdev=4.64) for nucleotide ORF1ab-CDS alignment, 54-97% (median=77.65, stdev=5.03) for ORF1ab-polyprotein alignment to 58-96% (median= 77.54, stdev=5.05) in nucleotide RdRp alignment (**Figure 3A-B**, **Supplementary tables 2-3**). The deltacoronavirus identified in *L. dominicanus* showed identities of 96.54%, 94.71% and 97.61% (**Supplementary table 2**) with the sequences from Gentoo penguin virus based on nucleotide RdRp, ORF1ab-CDS and ORF1ab-polyprotein alignments. Both viruses are closely related to those found primarily in Charadriiformes birds from distant geographical locations **(Supplementary figure 2).**

**Figure 3.**
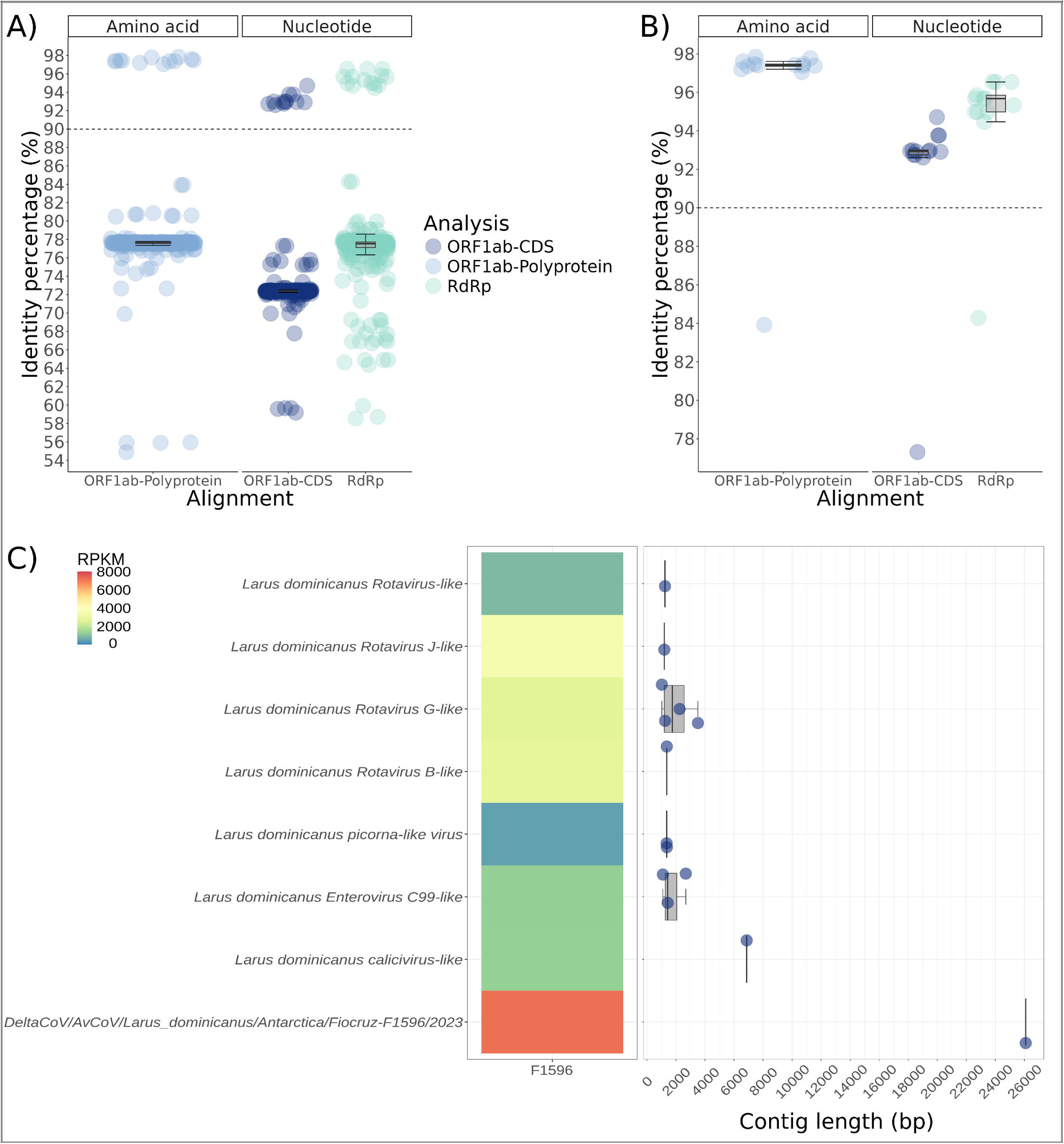
Genetic distances of Deltacoronavirus identified in *L. dominicanus* and those from NCBI, Reads per kilobase per million (RPKM) abundance and contig length of viruses identified. A) Distances among Deltacoronaviruses from clade depicted in Figure 2B. B) Distances among Deltacoronaviruses from clade depicted in Figure 2C. C) Heatmap showing the RPKM abundance and contig length of viruses identified in this study.

In addition to the Deltacoronavirus identified in this study, our analysis identified another 13 viral contigs showing best pairwise-alingment hits with four viral taxa (*Sedoreoviridae, Picornavirales, Picornaviridae*, *Caliciviridae,*) (**Figure 3C**). We identified 7 Rotaviruses sequences, 2 picorna-like virus sequences, 3 Enterovirus C99-like virus sequences and 1 calicivirus-like sequence. The coverM analysis showed the Deltacoronavirus as the most abundant virus, followed by the Rotaviruses (**Figure 3C**). The amino acid identities of these viruses in relation to their best hits ranged from 37-62% for Rotaviruses, 42-62% for picorna-like viruses, 94-97% for Enterovirus C99-like viruses, and 40% for calicivirus-like sequence (**Supplementary table 4**).

## DISCUSSION

Coronaviruses (CoVs) have a global distribution and can be found in various species of wild and domestic animals, serving as natural reservoirs hosts, virus evolution in these species allows the virus to accrue new genetic mutations through single amino acid changes and recombination events that may lead to the emergence of new serotypes or genera (Cui et al., 2019). Deltacoronaviruses are commonly found in wild birds and have been detected in all continents, including Antarctica (Chu et al., 2011; Wille and Holmes, 2020; Zamora et al., 2023). The deltacoronaviruses currently contain seven ratified species spread across three subgenera, all of which have been detected in wild birds (De-Groot et al., 2012; ICTV, 2022). Deltacoronaviruses appear to be the most prevalent in Anseriformes and Charadriiformes; however, their diversity in wild birds is likely much broader than currently documented. According to a recent review, avian coronaviruses have been detected in 15 orders, comprising 30 families, and across 108 species of wild birds (Wille and Holmes, 2020) across a wide range of host species, including gulls, shorebirds, penguins, passerines, and even bustards (Chamings et al., 2018; Chu et al., 2011; Wille and Holmes, 2020; Zamora et al., 2023).

The strain detected in our study was closely related with Gentoo penguin virus identified in *P. papua* from Antarctica, Isla Kopaitik, Base O’Higgin in 2014 but in locations 140 km apart, suggesting long term maintenance of this deltacoronavirus lineage in Antarctica and potential cross species transmission and infection of multiple avian species. According to ICTV (https://ictv.global/report_9th/RNApos/Nidovirales/Coronaviridae) viruses that share more than 90% aa sequence identity in the conserved replicase domains are considered to belong to the same species, thus the DeltaCoV identified belongs to a potential novel viral species that still not ratified by the ICTV, but that was tentatively named novel wild birds deltaCoV clade (Wille and Holmes, 2020). The same authors suggested that this Gentoo penguin virus could spillover from other birds such as skuas (*Stercorarus* spp.) or gulls inhabiting the same islands. Our results bring new evidence that support cross species transmission of this deltacoronavirus between species that inhabit the Antarctic environment. Although, the virus spread may be simultaneous taking place through interspecies spillover and penguins interpopulation migrations. *Larus dominicanus* distribution overlaps with that of gentoo penguins providing opportunities for inter species viral transmission (**Supplementary figure 2**). Moreover, this species has a generalist and opportunistic feeding behaviors, frequently scavenging and preying within penguin colonies (Barbieri, 2008; Favero and Silva, 1997). This carrion and eggs composition may facilitate *L. dominicanus* infection from infected penguin juveniles (Lebarbenchon et al., 2010). But, it remains to be evaluated if the DeltaCoV detected in *L- dominicanus* feces can actively infect *L. dominicanus* or it may be a result of excreta remains from Gentoo penguin infected juveniles.

There is currently no evidence that DCoV causes disease in wild birds (Chamings et al., 2018; Chu et al., 2011; Domanska-Blicharz et al., 2021, 2023). However, it is suggested that deltacoronaviruses may cause disease when introduced into poultry (Wille and Holmes, 2020) as described in farmed quail in Poland attributed to a Quail deltacoronavirus (Domanska-Blicharz et al., 2019). Deltacoronaviruses primarily infect birds, but their transmission to mammals is a relatively recent occurrence. Porcine deltacoronavirus (PDCoV; belonging to the Coronavirus HKU15 subgenus *Buldecovirus*) has emerged as a significant swine virus globally, originating from avian deltacoronaviruses (Woo et al., 2012). Initially identified in Asia in 2009, PDCoV outbreaks have been reported worldwide, causing severe diarrhea and vomiting in piglets (Kong et al., 2022; Woo et al., 2012; Zhang, 2016). Lately, human infections with PDCoV have also been documented, with the virus isolated from children suffering acute febrile illness (Lednicky et al., 2021). Experimental studies indicate that PDCoV can bind to both porcine and human APN receptors (Ji et al., 2022). Besides pigs, deltacoronaviruses have been detected in Asian leopard cats (*Prionailurus bengalensis*) and Chinese ferret badgers (*Melogale moschata*) in market settings (Dong et al., 2007). These findings highlight the growing concern about deltacoronaviruses’ potential to cross species barriers and pose risks to both animal and human health, emphasizing the importance of ongoing monitoring and surveillance efforts.

Approximately 40 species of sea birds breed on the Antarctic Peninsula, adjacent islands, and the Antarctic mainland (Shirihai, 2008). The Kelp Gull, widely distributed across the Southern Hemisphere, nests in South America, southern Africa, Australia, New Zealand, sub-Antarctic islands, and the Antarctic Peninsula, including Keller Peninsula on the southern side of the King George Island (Jaquim et al., 2009; Sander et al., 2006). During the austral winter, most Antarctic populations leave their breeding grounds and disperse across marine environments in the Southern Hemisphere, including Australia, New Zealand, South Africa, and South America (Krietsch, 2014). This migration provides an opportunity for the spread or emergence of new viruses, as the movement of *L. dominicanus* between regions could facilitate viral transmission across diverse ecosystems. In addition, all Laridae species live in colonies with high densities, where contact between infected individuals can easily occur and these factors may favor the infection of gulls with various coronaviruses (Arnal et al., 2015; Domanska-Blicharz et al., 2023).

We acknowledge the limitations of our study, as the fecal samples were collected from the environment rather than directly from the birds and could represent environmental contamination. However, most of the samples were collected short after the time of defecation. Additionally, the samples from the gull colony were taken in an area exclusively inhabited by gulls, with no penguin colonies present on Keller Peninsula. Penguins are only seen sporadically along the shoreline. Additionally, one would expect that environmental contamination with viral particles or viral genome would result in viral genomes sharded on many contigs with low coverage depth appearing in multiple fecal samples from the same colony. However, we only detected a DeltaCoV in one single sample and recovered its full genome with high genome coverage depth, which is indicative of high viral load being excreted by feces likely from an active infection. Therefore, we can confidently attribute the fecal samples to gulls rather than other bird species.

Influenza A viruses (IAVs) not only circulate in humans but also in domestic animals such as pigs, horses, and poultry, as well as in wild migratory birds, where more than 100 species of ducks, geese, swans, gulls, waders, and wild aquatic birds are considered natural reservoirs (Webster et al., 1992). Numerous studies have investigated the occurrence of IAVs in the avifauna of Antarctica, consistently showing that the local fauna is constantly exposed to IAVs infections (Austin and Webster, 1993; Barriga et al., 2016; de Souza Petersen et al., 2017; Hurt et al., 2016; Hurt et al., 2014; Ogrzewalska et al., 2022; Wallensten et al., 2006). However, all tested fecal samples were negative for influenza A viruses during monitoring carried out by our group in 2023, similarly to findings obtained in the same breeding season on the South Shetland Islands by (Munoz et al., 2024) and in 2022 by (Gomes et al., 2023). Therefore, these aggregated results suggest a low prevalence of IVA infection in Antarctic birds, corroborating literature data that have consistently observed a low prevalence (rates typically below 5%) of infection caused by IVA in Antarctic seabirds (Barriga et al., 2016; de Souza Petersen et al., 2017; Hurt et al., 2016; Hurt et al., 2014; Wille et al., 2020). Despite of that, in October 2023 (next breeding season after our study), the first case of Highly Pathogenic Avian Influenza (HPAI) was detected among the birds of the Antarctic region for the first time emphasizing the continuous monitoring and surveillance of influenza viruses among Antarctic fauna in warranted (Banyard et al., 2024).

Lastly, as metatranscriptomic approaches uncover the full breadth of viruses of any given sample (mostly viruses with RNA genome but also viruses with DNA genome) we detected and assembled full and partial virus genomes from four families known to infect other bird species. This data further supports that the Antarctica fauna harbors a diverse set of viruses that are evolving with their host for a long period of time (Smeele et al., 2018; Wille et al., 2020). Moreover, such rich genomic data will be key to characterize the current circulating viruses in Antarctica and the emergence of new virus introduction and their threats to wildlife.

## Conclusions

We successfully detected in feces of *L. dominicanus* and sequenced the complete genome of a novel *Buldecovirus*, closely related to a deltacoronavirus previously identified in *P. papua* from the Antarctic Peninsula. Although no Influenza A viruses were found, our analysis revealed additional viral contigs, which showed the highest sequence similarity to four viral taxa: *Sedoreoviridae, Picornavirales, Picornaviridae,* and *Caliciviridae.* This underscores the vast, largely unexplored viral diversity in Antarctic ecosystems. Given the limited and fragmented understanding of the ecology of Influenza A viruses and coronaviruses in these remote environments, our findings emphasize the critical need for ongoing surveillance and research. Such efforts are vital to elucidate the dynamics of IAV circulation in Antarctic wildlife, ensuring the protection of both animal and human health.

## CONFLICTS OF INTEREST

The author(s) declare that there are no conflicts of interest.

## FUNDING INFORMATION

This study was partly supported by INOVA – EDITAL 2/2018 - GERAÇÃO DE CONHECIMENTO [grant number: 4720463444], and National Council of Technological and Scientific Development (CNPq)/MCTIC/CAPES/FNDCT N° 21/2018 – PROANTAR [grant number: CNPQ 442646/2018-6]. Fundação para o Desenvolvimento Científico e Tecnológico em Saúde- FIOTEC [process: IOC-026-FIO-21] in award a scholarship.

## ACKNOWLEDGMENTS

Permissions to collect samples were granted by the Environmental Assessment Group of the Brazilian Antarctic Program (GAAm-PROANTAR XLI) for specific locations (Antarctic Specially Protected Areas, ASPAs) within the Antarctic region. These locations include the Western Shore of Admiralty Bay (ASPA 128), Lions Rump (ASPA 151), Potter Peninsula (ASPA 132), Ardley Island (ASPA 150), Byers Peninsula (ASPA 126), Harmony Point (ASPA 133), and Deception Island (ASPA 140). No permission was required for sampling in other areas as they did not have any access restrictions or specific regulations in place.

FIOANTAR Working Group (https://fioantar.fiocruz.br/equipe), the Brazilian Antarctic Program – PROANTAR and Vice-Presidency of Production and Innovation in Health (VPPIS). We acknowledge the Norwegian Polar Institute’s Quantarctica package. GLW is supported by the Conselho Nacional de Desenvolvimento Científico e Tecnológico (CNPq) through their productivity research fellowships (307209/2023-7).

---

**Supplementary figure 2 –** Distributions of *Larus dominicanus*, *Pygoscelis papua* and bird species in which coronaviruses closely related to ours were detected, using data from the BirdLife database (BirdLife, 2023). The map made in QGIS 3.32.2 (QGIS, 2024).

**Supplementary table 1 -** Localization of the collection sites in the present study and the number of bird fecal samples collected, South Shetland Islands, January-February 2023. ASPA - Antarctica Specially Protected Area.

**Supplementary table 2 -** Distance analysis of Deltacoronavirus from *L. dominicanus* and deltacoronaviruses from GenBank (attached separately).

**Supplementary table 3 -** Statistics from distance analysis.

**Supplementary table 4 -** The amino acid identities of detected viruses in relation to their best hits

## Notes

### Competing Interest Statement

The authors have declared no competing interest.

