## Supplementary figures and images for "Detection and full genome sequencing of a Deltacoronavirus and other bird associated viruses from feces of the kelp gull (*Larus dominicanus*) sampled at the South Shetland Islands Antarctica"

### Supplementary figure 2

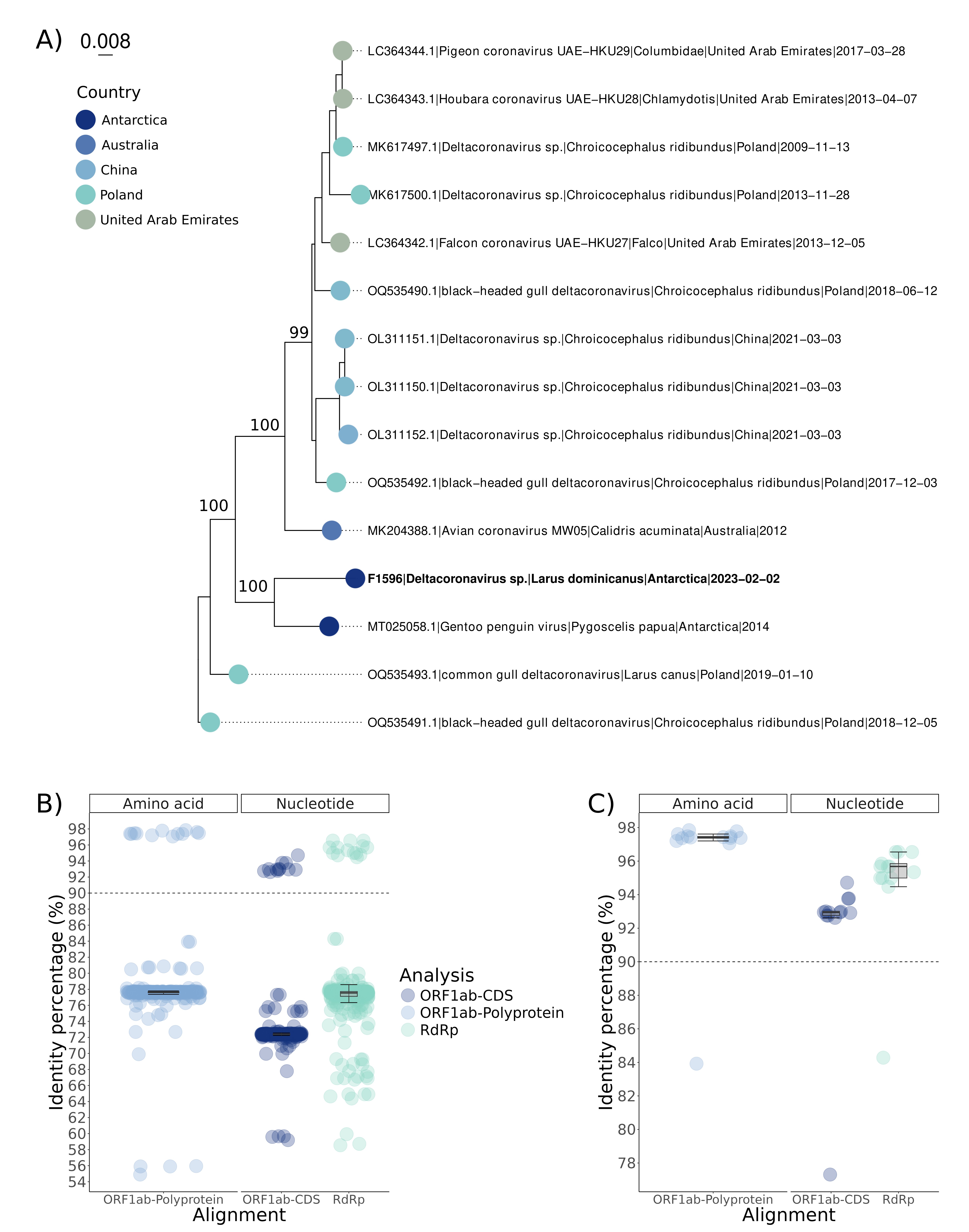

### Supplementary figure 2

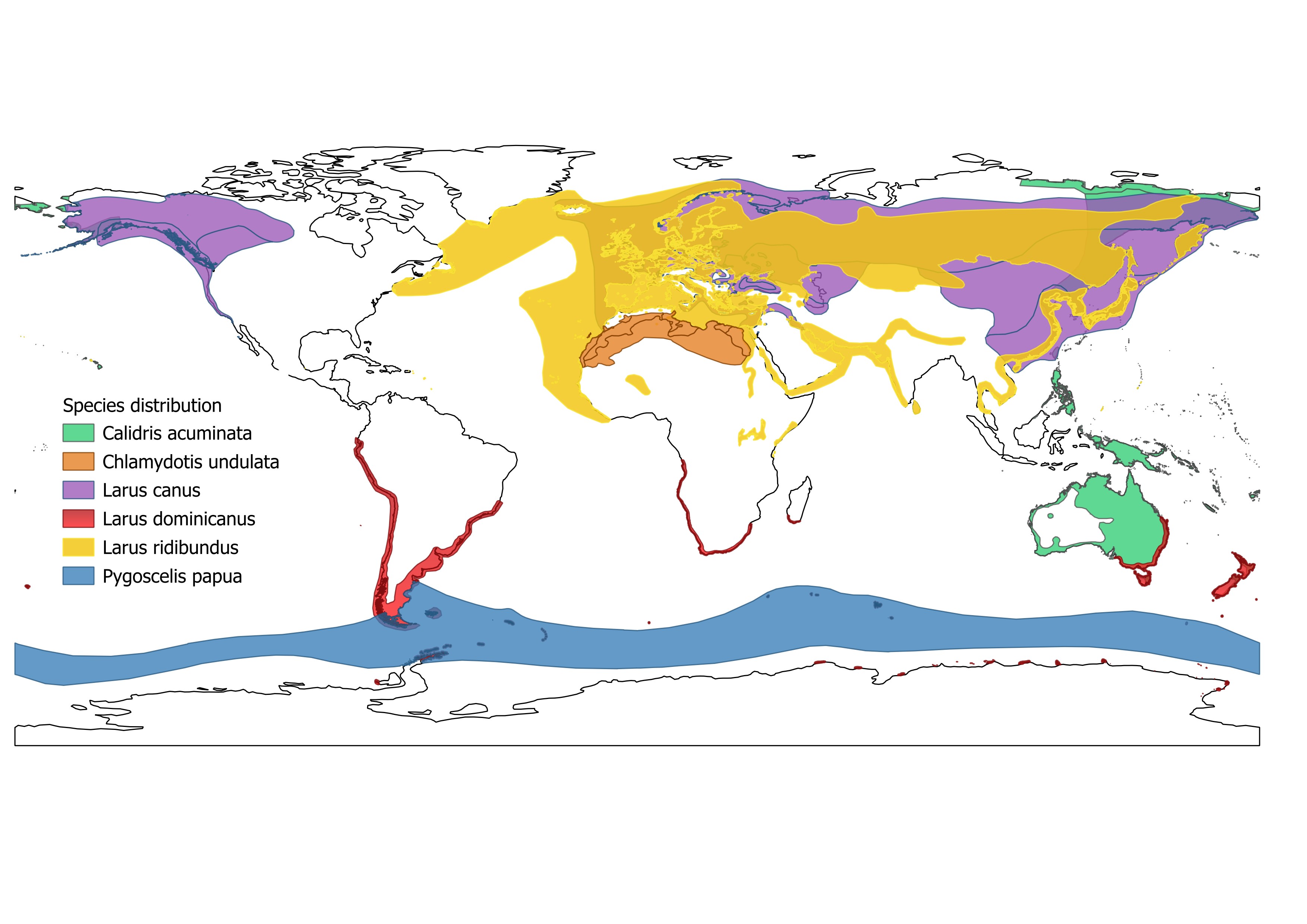
